# Demyelination induces transcriptional reprograming in proprioceptive and Aβ rapidly adapting low-threshold-mechanoreceptor neurons

**DOI:** 10.1101/2021.11.23.469748

**Authors:** Benayahu Elbaz, Lite Yang, Braesen Lee Rader, Riki Kawaguchi, Maria Traka, Clifford J Woolf, William Renthal, Brian Popko

## Abstract

Schwann cells, the main glial cell in the peripheral nervous system (PNS), ensheath bundles of small unmyelinated axons or form myelin on larger axons. PNS injuries initiate transcriptional reprograming in both Schwann cells and sensory neurons that promotes regeneration. While the factors that initiate the transcriptional reprograming in Schwann cells are well characterized, the full range of stimuli that initiate this reprograming in sensory neurons remain elusive. Here, using a genetic model of Schwann cell ablation, we find that Schwann cell loss results in transient PNS demyelination without overt axonal loss. By profiling sensory ganglia at single-cell resolution we show that this demyelination induces transcriptional reprogramming preferably in proprioceptive and Aβ RA-LTMR neurons. Transcriptional reprograming is assumed to be a cell autonomous response of sensory neurons to mechanical axonal injury. By identifying similar reprograming in non-injured, demyelinated neurons, our study suggests that this reprograming represents a non-cell autonomous transcriptional response of sensory neurons to the loss of axon-Schwann cell interactions.

**Highlights:**

1. Ablation of Schwann cells results in transient PNS demyelination, without overt axonal loss.
2. Schwann cell loss results in transcriptional reprograming in specific sensory neurons.
3. Spinal nerve transection (mechanical injury of axons) and demyelination (intact axons) induces similar transcriptional responses in DRG neurons.
4. The transcriptional response to demyelination among DRG neurons is specific to the large myelinated proprioceptive and Aβ RA-LTMR neurons.

## Introduction

Peripheral nervous system (PNS) myelin facilitates rapid nerve conduction velocities and provides trophic support to axons. During development, neural crest derived Schwann cell precursor cells comigrate with the growing axons to their terminal destinations, where they differentiate to myelinating or non-myelinating Schwann cells (Jessen et al., 2015). The peripheral axons provide trophic and mitogenic factors to the Schwann cells, while Schwann cell support the long-term survival of the axons and the formation and organization of nodes of Ranvier (Corfas et al., 2004). Non-myelinating Schwann cells engulf low (<1 μm) caliber axons. High levels of neuregulin 1 type III expressed on the surface of high caliber (>1 μm) axons triggers the Schwann cell myelination program resulting in a one-to-one ratio between myelinating Schwann cells and axons (Michailov et al., 2004; Taveggia et al., 2005). Mutations in Schwann cell specific genes, such as Connexin-32 (Cx32) and Myelin Protein Zero (Mpz), that affect normal Schwan cell function, result in demyelination and axonal degeneration (Berger et al., 2006). In addition, impaired Schwann cell function and demyelination accompanied by axon damage is a debilitating result of chemotherapy treatment (Imai et al., 2017) and of both type 1 and type 2 diabetes (Gonçalves et al., 2017). Demyelination of the PNS dramatically affects sensory neurons and leads to loss of sensation, and in some patients, debilitating neuropathic pain. Strikingly, despite the clinical importance, the effect of PNS demyelination on sensory neuron gene expression is largely unknown.

PNS injury initiates in both Schwann cells and sensory neurons transcriptional reprograming that promotes in both cell types, a new transcriptional state that promotes regeneration. Schwann cells that lose their axon-glia interaction upon injury dedifferentiate and transform into repair Schwann cells that aid axonal regeneration. This transcriptional reprograming is controlled mainly by the early transcription factors c-Jun (Arthur-Farraj et al., 2012), the transcription factor STAT3 (Benito et al., 2017), by chromatin modifications (Ma and Svaren, 2018) and by Merlin (Mindos et al., 2017). In parallel, sensory neurons responds to mechanical axonal injury by a transcriptional reprograming that suppresses sensory neuron cell identity and promotes neuronal regeneration. In sensory neurons, this reprograming depends on the early transcription factor ATF3 (Renthal et al., 2020). Upon completion of neuronal regeneration and reinnervation, the original cell identity is reestablished, and Schwann cell axon-glia interactions are restored (Jessen and Mirsky, 2016, 2019).

The signals that initiate the transcriptional reprograming in sensory neurons upon injury are poorly understood. Signals from the injured axon to its counterpart Schwann cell instruct the initial response of Schwann cells to axonal injury (Catenaccio et al., 2017; Vaquié et al., 2019). Nevertheless, it is unknown exactly what triggers the injury induced transcriptional reprograming in sensory neurons. Previous studies suggested that this transcriptional reprograming is a cell autonomous property of sensory neurons, activated solely by mechanical injury of the neurons (Cheng et al., 2021; Cho et al., 2013). In these studies, transcriptional reprograming in sensory neurons was documented in spinal nerve transection, sciatic nerve transection, and sciatic nerve crush, and was not observed in non-mechanical models that promote chronic pain, such as models of inflammatory and chemotherapy-induced pain, where only a few genes were activated in sensory neurons (Renthal et al., 2020). Nevertheless, it is not clear if Schwann cell axon-glia interactions are impaired in these non-mechanical models of PNS injury, and the role of Schwann cells axon-glia interactions in the transcriptional state of sensory neurons remain elusive.

To study if Schwann cells have a specific role in modulating the transcriptional state of sensory neurons, we characterized a genetic mouse model of Schwann cell ablation. We find that Schwann cell loss results in a transient PNS demyelination without overt axonal loss. Using a combination of bulk and single-cell seq we find that this demyelination and neuropathy results in transcriptional reprograming in sensory neurons. This transcriptional response was very similar to the transcriptional response of sensory neurons to mechanical injury, suggesting that loss of axon-glia interactions contributes to the transcriptional reprograming in sensory neurons. Nevertheless, while mechanical injury affects nearly all the sensory neurons with injured axons, we find that the transcriptional response to Schwann cell loss occurs preferentially in heavily myelinated proprioceptive neurons and Aβ rapidly adapting low-threshold-mechanoreceptor (Aβ RA-LTMR) neurons, suggesting that it is demyelination that initiates the transcriptional reprograming in these sensory neurons, in our model.

## Results

### 1. Genetic ablation of Schwann cells results in neuropathy and demyelination of the PNS without overt axonal loss

For our studies we focused on the well characterized PLP-CreER^T^;ROSA26-eGFP-DTA (DTA) model, in which the expression of the diphtheria toxin A-subunit (DT-A) is prevented by upstream “lox-stop-lox” (LSL) cassette, and only upon tamoxifen-induced PLP-CreER^T^ mediated recombination, the stop sequence is removed, and the expression of DT-A is released, resulting in cell loss. CNS demyelination in DTA mice has been characterized in detail (Traka, 2019; Traka et al., 2010, 2016). Nevertheless, the PLP-CreER^T^ line drives recombination in Schwann cells as well (Doerflinger et al., 2003), suggesting possible PNS demyelination in tamoxifen treated animals. About 80% of the myelinated axons of the sciatic nerve are sensory axons (Schmalbruch, 1986). Based on this, we focused on demyelination of the sciatic nerve and on the transcriptional response of the sensory neurons in the lumbar DRGs (L3-L5) that project through the sciatic nerve (Rigaud et al., 2008). Our phenotypic characterization revealed that in DTA mice the conduction velocity in the sciatic nerve is substantially reduced 21 days post tamoxifen administration (PID21), and full functional recovery is achieved at PID42 (Fig. 1A). Interestingly, the clinical score of these mice peaks at PID21, before the onset of the reported CNS pathology, and reduces at PID28, in parallel to the functional recovery in the PNS (Fig. 1B). While complete functional recovery of the PNS is achieved at PID42 (Fig. 1A), the clinical score dramatically increases at this time point (Fig. 1B), due to widespread CNS demyelination.

**Figure 1:**
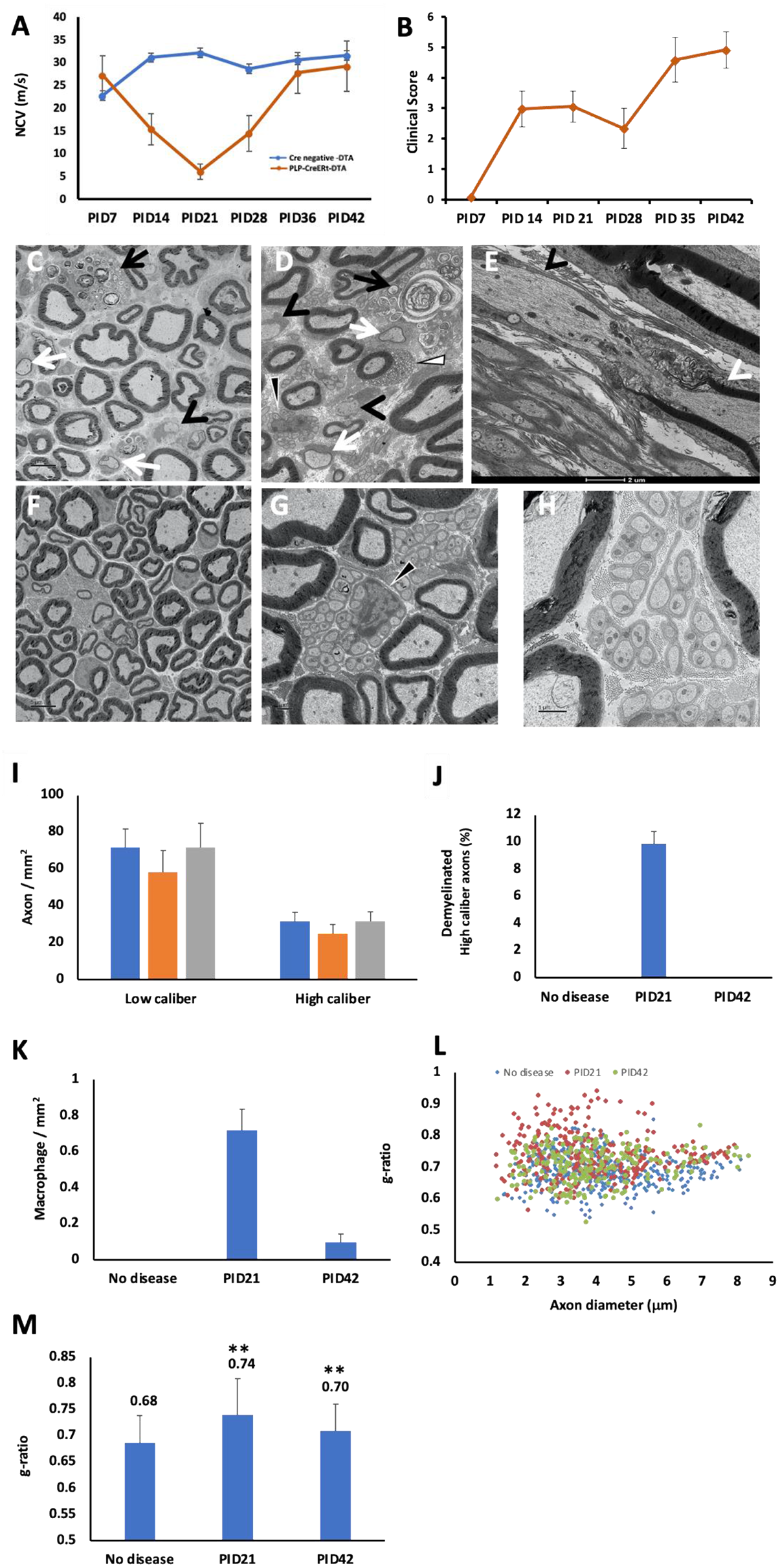
Schwann cells ablation results in demyelination without axonal loss. (A) Conduction velocity in the sciatic nerve at days 7-42 post tamoxifen injection (PID). (B) Clinical scores of the demyelinated mice. Clinical scores were based on a scoring system developed for phenotypic characterization of demyelination in this line (Traka, 2019; Traka et al., 2010, 2016). (C-E) EM of the sciatic nerve at PID21: demyelinated large caliber axons (black arrowheads), remyelinated large caliber axons (white arrows), phagocytic macrophages (black arrows), apoptotic myelinating Schwann cells (D, white triangle) and Remak bundles that are not fully engulfed by basal lamina (D, black triangle). (E) Segmented demyelination-on the same axon the left internode was absent (black arrowhead) while the right internode was present (white arrowhead). (F-H) At PID42, the large caliber axons were remyelinated. (G-H) Remak bundles that are not fully engulfed by basal lamina at PID42. (I) No axonal loss was observed. (J) About 10% of the high caliber axons (diameter >1 m) were demyelinated at PID21. (K) Quantification of phagocytic macrophages in the Sciatic nerve. (L) Reduced myelin thickness at PID21 (red) and PID42 (green) compared to control (blue). (M) Myelin thickness, quantified results. For panels A and B PLP CreERt;ROSA26-eGFP-DTA mice; n=14. For the Cre negative ROSA26-eGFP-DTA mice; n=11. For panels C-M n=3 for no disease, n=4 for PID21, and 3 for PID42.

Based on these electrophysiological results, we focused on the PID21 and PID42 time points, and assessed the morphology of the sciatic nerve by EM. We found that at PID21 the sciatic nerve is characterized by appearance of demyelinated and remyelinated large caliber axons, aberrant morphology of Remak bundles, infiltration of phagocytic macrophages, and reduced myelin thickness (increased g-ratio) (Fig. 1). In addition, we observed segmented demyelination in which on the same axon one internode was absent, due to loss of the myelinating Schwann cell, and the adjacent internode was still present (Fig. 1E). This segmented demyelination suggests that even for the axons that seem myelinated in cross section, the axons are likely demyelinated at one or more points along the long axon, which results in the observed severe drop in nerve conduction. We did not detect axonal loss (Fig. 1I), nor did we observe the Büngner bands that characterize loss of myelinated axons or collagen pockets that characterize loss of nonmyelinated fibers. At PID42, all large caliber axons were well myelinated (Fig. 1F), but an aberrant morphology of Remak bundles that are not fully engulfed by basal lamina was still observed (Fig. 1G-H).

### 2. Demyelination of the PNS induces expression of RAG genes in lumber DRG neurons

Based on the morphological changes in both myelinated fibers and Remak bundles in the sciatic nerve detected in the PLP-DTA mouse, we hypothesized that the Schwann cell loss might result in changes in gene expression in the sensory neurons in the lumber DRGs that project through the sciatic nerve. Based on the electrophysiology and EM studies, we decided to focus our transcriptomic studies on PID21 and PID42, and harvested the sciatic nerve and the lumbar DRGs (L3-L5) for bulk RNA-seq. In the DRGs, we found that at peak disease (PID21), expression of the known key injury-induced Regeneration Associated Genes (RAGs) *Atf3, Sox11* and *Sprr1a*, is induced (Fig. 2A). In addition, transcripts involved in inflammation (such as *Trem2, Stab1* and *Tgfb1*) were also induced (Fig 2C). This may reflect similar infiltration of macrophages into the DRGs and an inflammatory response in the cells in the DRG that occurs after nerve injury. Transcripts involved in oxidative phosphorylation and ATP production were down regulated in the DRGs (Fig. 2B), suggesting that the cells in the DRGs suffer from mitochondrial damage. Nevertheless, this may also reflect death of DRG resident Schwann cells. In the sciatic nerve, transcripts involved in cell mitosis and cell division were down regulated at the peak of demyelination (Fig. 2D and E), and transcript involved in developmental processes were upregulated (Fig. 2F). This probably reflects the effect of concurrent demyelination and remyelination observed at the peak of demyelination in the sciatic nerves (Figure 1). The induction of RAGs *Atf3, Sox11* and *Sprr1a*, was specific for the DRG and was not observed in bulk RNA-seq of the sciatic nerve, suggesting that the induction of RAGs is specific to the cells in the DRGs. To directly assess if the induction of RAGs is specific to sensory neurons, we analyzed the transcriptome of DRG neurons directly by Single Nucleus RNA-seq (snRNAseq).

**Figure 2:**
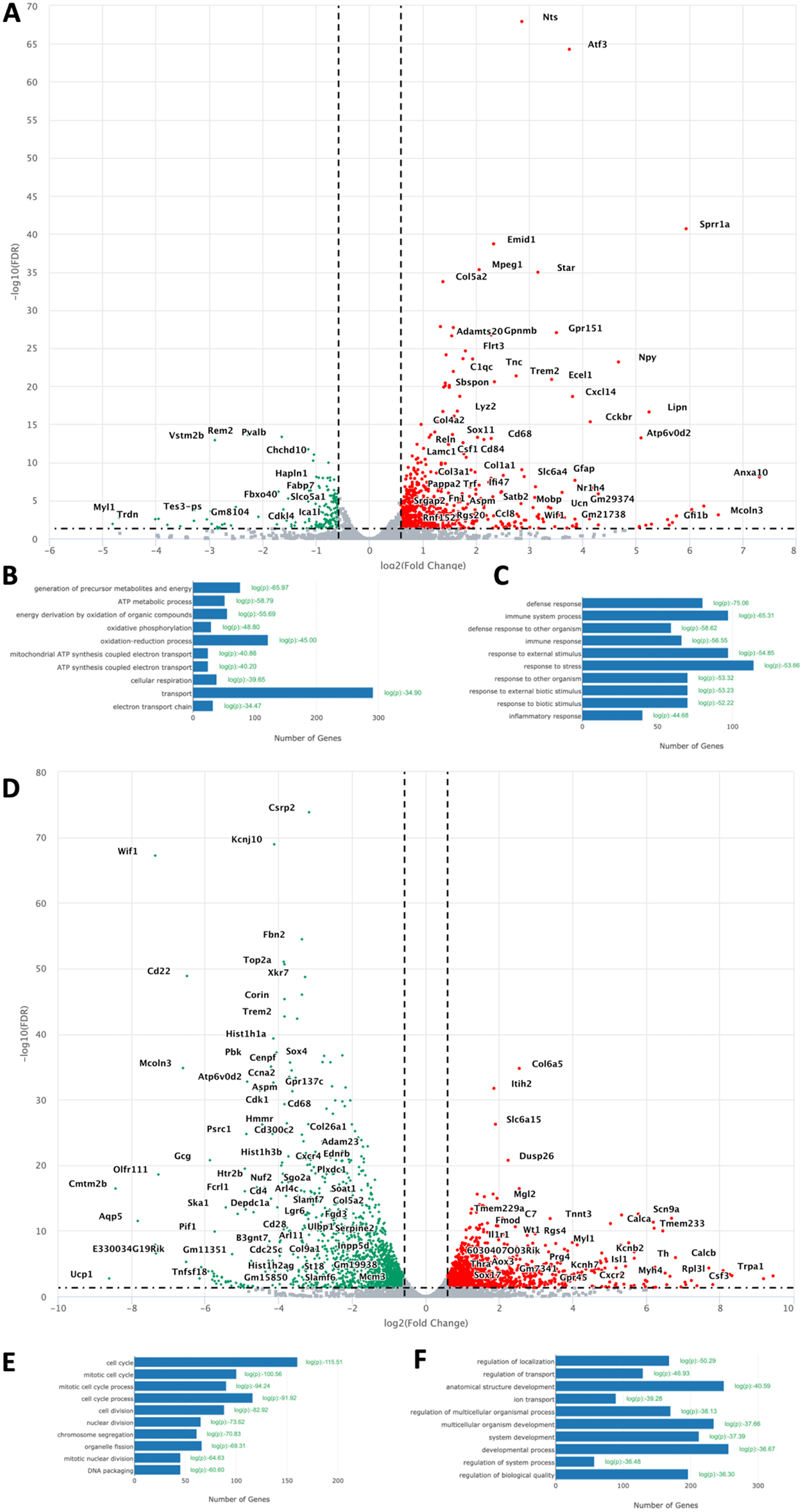
Demyelination induces the expression of Regeneration Associated Genes in DRG neurons. Eight-week-old Plp-CreERt;ROSA26-eGFP-DTA mice and Cre negative ROSA26-eGFP-DTA mice (used as control) were injected with tamoxifen, and L3-L5 DRGs were dissected for bulk RNA-seq. (A) Volcano plot of the Differentially Expressed Genes (DEG) in the DRG at PID21. Green-downregulated genes, red-upregulated genes. (B) Biological processes analysis of downregulated genes. (C) Biological processes analysis of upregulated genes. (D) Volcano plot of the Differentially Expressed Genes (DEG) in the sciatic nerve at PID21. Green-downregulated genes, red-upregulated genes. (E) Biological processes analysis of downregulated genes. (F) Biological processes analysis of upregulated genes. n = 2 biologically independent experiments. (FDR ≤ 0.05, fold-change ≥ 2).

### 3. New Injured Transcriptional State in DRG Neurons After Schwann cells loss

To characterize the transcriptional changes in sensory neurons in response to Schwann cell loss and demyelination at the single-cell level, we sequenced DRG nuclei from naive mice (no disease), from mice at the peak of demyelination (PID21) and from mice following PNS remyelination (PID42). For these studies we harvested the cell bodies of the sensory neurons that project through the sciatic nerve (L3-L5 DRGs). Using a method that enriches for neuronal nuclei (Yang et al., 2021), we profiled 14,065 nuclei that passed quality control. The number of cells analyzed in each condition is provided in Tables S2-S4. To classify cell types, DRG nuclei from naive and demyelinated DRGs were initially clustered together based on their gene expression patterns. Dimensionality reduction (t-distributed stochastic neighbor embedding [tSNE]) revealed distinct clusters of cells, with neuronal clusters expressing *Rbfox3*, and non-neuronal clusters expressing *Sparc* (Fig. S1). We identified nine naive cell types including two clusters of glial cells (725 nuclei) and six different subtypes of sensory neurons (12,726 nuclei). We observed six naive DRG neuron subtypes: *Fam19a4+/Th+* C-fiber LTMRs (cLTMR1), *Tac1+/Gpx3+* peptidergic nociceptors (PEP1), *Tac1+/Hpca+* peptidergic nociceptors (PEP2), *Mrgprd+* non-peptidergic nociceptors (NP), *Nefh+* A fibers including Aβ low-threshold mechanoreceptors (Aβ-LTMRs) and *Pvalb+* proprioceptors (NF1, NF2) and *Sst+* pruriceptors (SST). We also observed non-neuronal cell types, including *Apoe+* satellite glia and *Mpz+* Schwann cells (Fig. 3A, and Table S1). Moreover, we identified an additional cluster of nuclei (614 nuclei, ∼4% total) that are derived mostly from mice at the peak of PNS demyelination (86.8%). Nuclei in this cluster express high levels of the known peripheral nerve injury-induced genes *Atf3, Sox11* and *Sprr1a*, such that they were defined as “injured” (Renthal et al., 2020) (Fig. 3A). In line with previous studies (Nguyen et al., 2019; Renthal et al., 2020), we found that 1.3% of the nuclei at “no disease” were injured, which may represent routinely occurring fight wound injuries in group-housed mice. The percentage of the injured nuclei increased to 7.9% upon demyelination and was reduced to 3.4% upon remyelination (Fig 3B). The DRG cell types identified here express unique patterns of ion channels, G protein-coupled receptors (GPCRs), neuropeptides, and transcription factors that are consistent with previous studies (Fig. 3C) (Renthal et al., 2020; Sharma et al., 2020; Usoskin et al., 2015; Zeisel et al., 2018; Zheng et al., 2019). Comparison between the neuronal injury-induced genes in spinal nerve transection (Renthal et al., 2020) and the marker genes in the “injured” cluster observed here upon demyelination revealed that the two “injured” transcriptional states are very similar (Fig. 3D).

**Figure 3:**
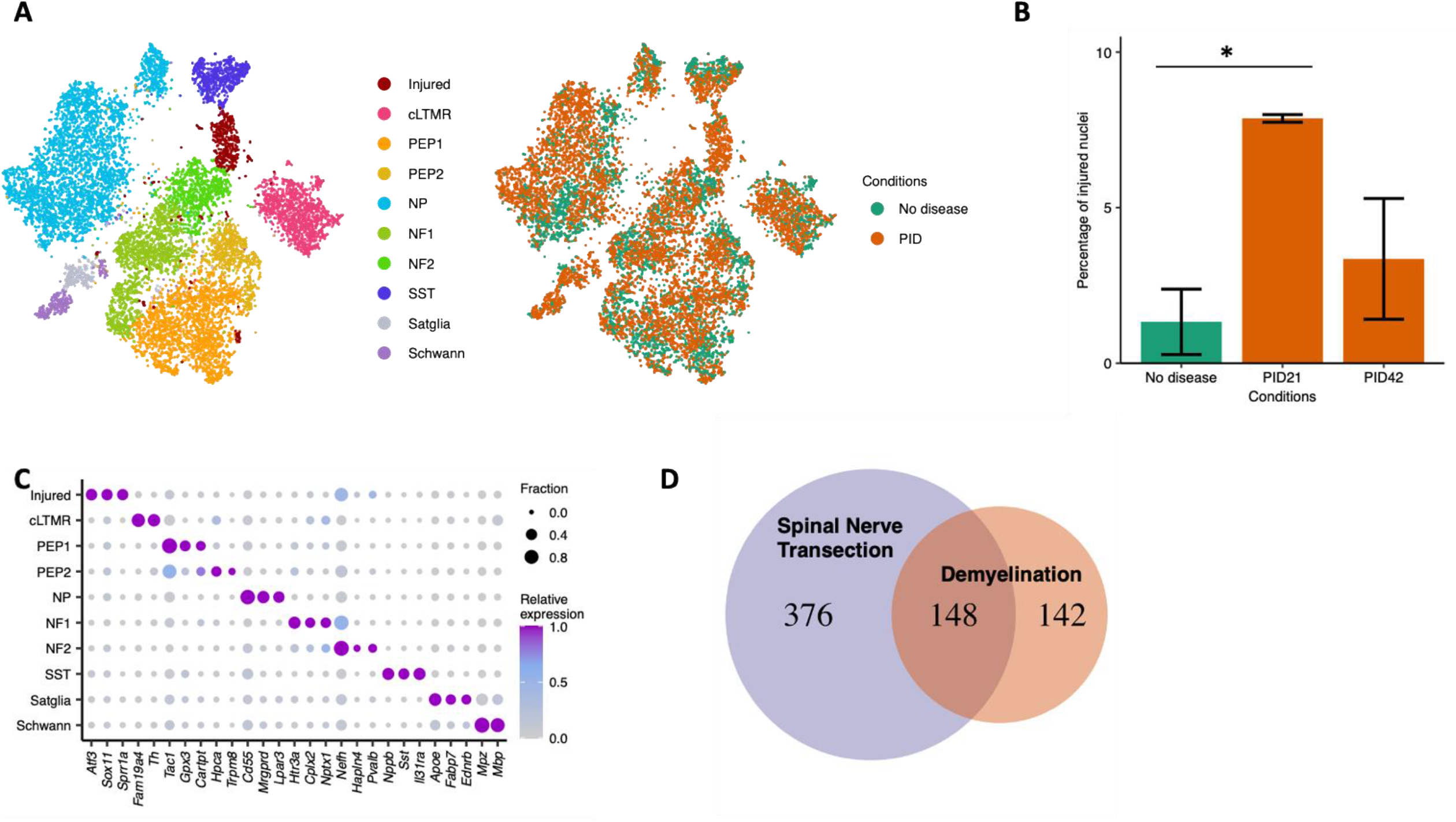
New Injured Transcriptional State in DRG Neurons After Schwann cells loss. (A). Eight-week-old Plp-CreERt;ROSA26-eGFP-DTA mice and Cre negative ROSA26-eGFP-DTA mice (used as control) were injected with tamoxifen, and L3-L5 DRGs were dissected for snRNA-seq studies at PID21 and PID42. tSNE plot of 14,065 nuclei from Cre negative ROSA26-eGFP-DTA mice (No disease) and Plp-CreERt;ROSA26-eGFP-DTA mice at PID21 and PID42. Nine naive cell types, including glial cells and different subtypes of sensory neurons, and one cluster of injured nuclei were identified. Nuclei are colored by cell type (left) or condition (right). (B). Bat plot showing average percentage of nuclei that are in the injured cluster across duplicates in each condition (n=2). Error bars denote standard deviations. Two-way ANOVA was conducted to compare the percentage across conditions. F(2, 3) = 13.74, p = 0.03. PID21 shows significant increase in injured nuclei compared to no disease control (Bonferroni post hoc, p = 0.03). (C). Dotplot displaying the relative gene expression of marker genes (columns) in the injured cluster and each naive cell type (rows). The fraction of nuclei expressing a marker gene is calculated as the number of nuclei in each cell type that express a gene (>0 counts) divided by the total number of nuclei in the respective cell type. Relative expression in each cell type is calculated as mean expression of a gene relative to the highest mean expression of that gene across all cell types. (D). Overlap between the common neuronal injury-induced genes in spinal nerve transection (see methods) and marker genes in the injured cluster (log2FC > 0.25, P < 0.05, comparing nuclei from the injured cluster to all other nuclei) (hypergeometric test, p=2e-102).

### 4. Schwann cell loss induces transcriptional reprograming selectively in proprioceptive and Aβ rapidly adapting low-threshold-mechanoreceptor neurons

The sensory neurons in the DRGs represent a heterogenous population of neurons that vary in their degree of myelination. We therefore hypothesized that unlike mechanical injury, demyelination may have a unique impact specifically on heavily myelinated sensory neurons (A-fibers) and little to no effect on non-myelinated sensory neurons (C-fibers). To test this hypothesis, we investigated whether the gene expression program induced by demyelination differs between distinct DRG neuronal subtypes. To classify the cell types of nuclei in the injured cluster whose cell-type-specific marker genes are down regulated, we co-clustered the injured nuclei identified in our study to those previously classified in Renthal et al. (Renthal et al., 2020) (Fig. 4A). To validate the cell type assignment, we anchored the injured nuclei to the naive nuclei in our study whose cell identity has been identified previously. Cell type assignments from the two approaches had 85.7% concordance (Figure S3). Consistent with prior observations (Nguyen et al., 2019; Renthal et al., 2020), a small fraction of the NF1 and NF2 cells (0.18% and 0%, respectively) display an injured transcriptional state in non-diseased mice, but 8.1% of the cells in the NF1 cluster and 40.1% of those in the NF2 cluster exhibited the injured state at the peak of demyelination (PID21). The percentage of injured NF1 and NF2 nuclei declines to 1.3% and 5.3% at the remyelination (PID42), respectively (Fig. 4B). This data suggests that the transcriptional response to demyelination among DRG neurons is preferentially induced in the neurons that compose clusters NF1 and NF2, which are the heavily myelinated A fibers including *Nefh+* Aβ low-threshold mechanoreceptors (Aβ-LTMRs) and *Pvalb+* proprioceptors. To validate these results by immunohistochemistry, we used the marker Neurofilament Heavy Polypeptide (NEFH) that marks Aβ-LTMRs and proprioceptors neurons in the DRG (Renthal et al., 2020; Usoskin et al., 2015). In line with the snRNA-seq results, the expression of ATF3 is strongly induced at the time of peak disease (PID21), specifically in large diameter A fibers marked by NEFH (Fig. 4B and C). Analysis of the differentially expressed genes in the NF2 cluster upon demyelination revealed that the demyelination induced transcriptional changes in this cluster representing a transcriptional reprograming, very similar to the one we observed in sensory neurons upon mechanical transection or crush injury (Renthal et al., 2020) (Fig 4E). The transcripts induced by demyelination and their similarity to those that occur after spinal nerve transection are listed in Table S5.

**Figure 4:**
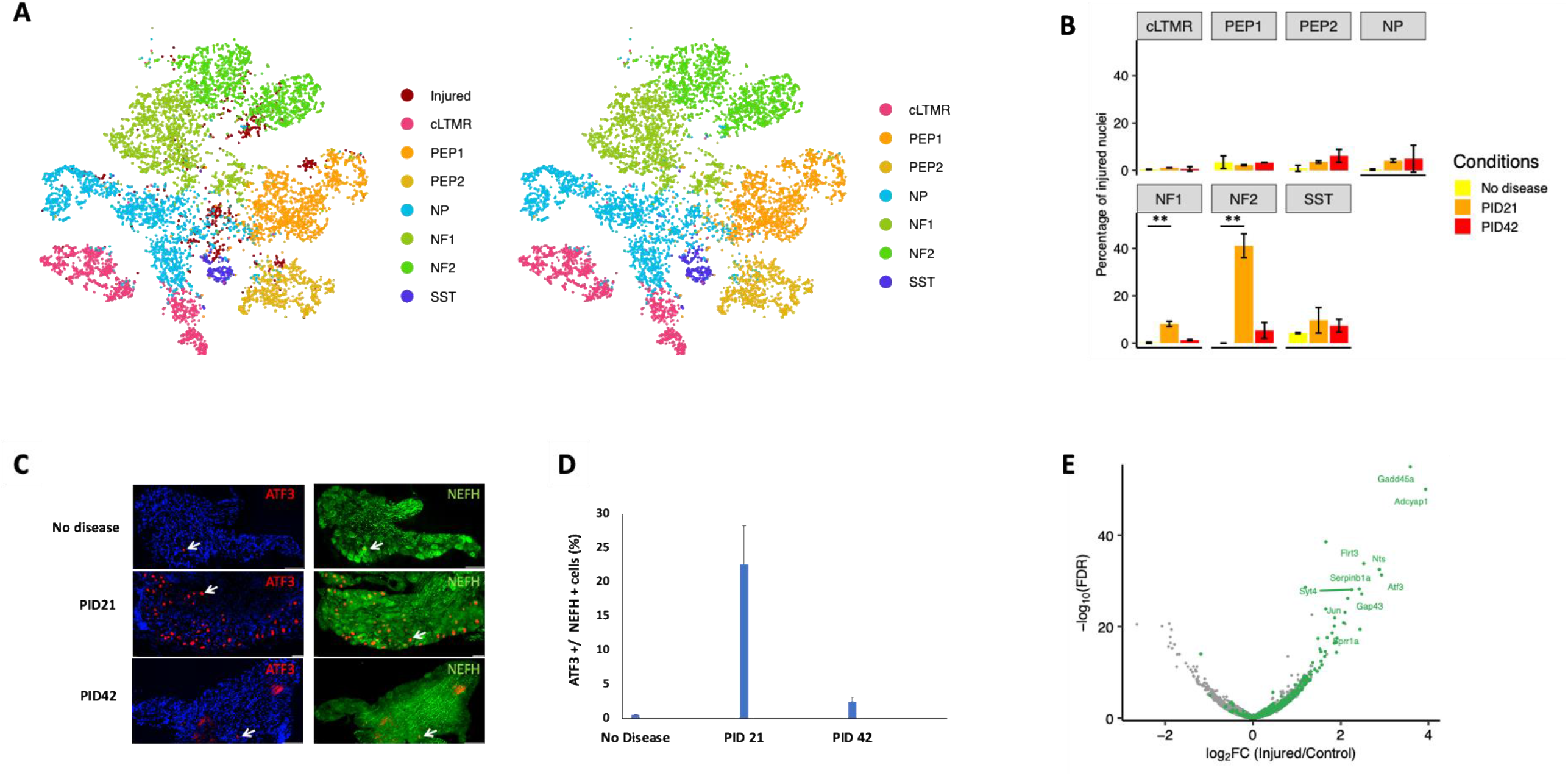
Schwann cell loss induced transcriptional reprograming selectively in proprioceptive and Aβ rapidly adapting low-threshold-mechanoreceptor neurons. (A) Classification of injured DRG nuclei after demyelination by co-clustering with injured nuclei after Spinal Nerve Transection. Nuclei after demyelination were in dark red, and nuclei after Spinal Nerve Transection were colored by their respective cell types (left). Nuclei after demyelination were annotated by projecting the Spinal Nerve Transection neuronal subtype classification onto the cluster. All nuclei were colored by their respective cell types (Right). (B). Bar plot showing average percentage of nuclei that are in the injured cluster for each subtype in each condition. Error bar shows the standard deviation across biological replicates. ANOVA was conducted to compare the average percentage of injured nuclei across conditions in each subtype. NF1: F(2, 3) = 87.68, p = 0.002; NF2: F(2, 3) = 81.53, p = 0.002. PID21 shows an increase in percentage of injured nuclei compared to no disease in both NF1 and NF2 (Bonferroni post hoc, p < 0.01). (C-D). Verification of the results by IHC; (C) the expression of ATF3 is induced at peak disease (PID21) specifically in Aβ-LTMR and Proprioceptor neurons, marked by NEFH; (D) Quantitative analysis. (E). Volcano plot displays differentially expressed genes between injured NF2 nuclei from PID21 and nuclei from no disease. Common-injury induced genes in Spinal Nerve Transection are colored green. Top 10 most differentially expressed genes that are also common-injury induced genes in Spinal Nerve Transection are labeled.

## Discussion

Mechanical injury of peripheral nerve axons leads to Wallerian degeneration followed by a regrowth of axons toward their targets and their reinnervation. This process is tightly regulated by a cascade of transcriptional events that result in the conversion of mature sensory neurons into actively growing cells (Abe and Cavalli, 2008; Chandran et al., 2016). In this study we used a genetic model of Schwann cell ablation to study the effect of PNS demyelination on the transcriptome of sensory neurons, without the confounding effect of neuronal/axonal injury. We found that ablation of Schwann cells results in a transient PNS demyelination, in which about ten percent of large caliber (>1 μm) axons are demyelinated, as observed in similar models of induced Schwann cell loss (Gerber et al., 2019). We observed that this peripheral demyelination results in a profound transcriptional reprogramming of a distinct set of sensory neurons, including the induction of RAGs and the downregulation of genes that define transcriptional identity. We found that this transcriptional reprogramming is transient and reversible, as the transcriptional states of injured neuronal nuclei return to their naive states within weeks (Fig. 4), coinciding with remyelination (Fig. 1).

Previous studies revealed a core transcriptional reprograming in sensory neurons in response to various peripheral nerve mechanical injuries (Renthal et al., 2020). We found a very similar transcriptional reprogramming in sensory neurons upon demyelination (Fig. 3 and 4). The similar transcriptional reprograming in both conditions suggests that this pattern of reprograming may represent a non-cell autonomous transcriptional response of sensory neurons to the loss of axon-Schwann cell interactions rather than simply being a reaction to axonal damage. A similar phenomenon was recently observed following optic nerve injury, where the response of retinal ganglion cells to optic nerve injury turns out to be dictated by amacrine cells (Sergeeva et al., 2021; Zhang et al., 2019). However, while mechanical axonal injury produces transcriptional changes in effectively all the injured sensory neurons, our studies here reveal a neuronal sub-type vulnerability to PNS demyelination of Aβ RA-LTMR and proprioceptor neurons, which have the highest axon caliber and thickest myelin, suggesting that the transcriptional reprograming in these neurons can be evoked by demyelination without direct physical axonal injury. Interestingly, despite the marked aberrant phenotype in non-myelinating Schwann cells following the Schwann cell ablation (Fig. 1), we did not observe substantial changes in the transcriptome of C-fiber neurons (Fig. 4). A similar neuronal specificity is present in Charcot-Marie-Tooth disease type 1A (CMT1A) patients suffering from PNS demyelination. In these patients, despite the presence of abnormal non-myelinating Schwann cells, no effect on the unmyelinated axons was noted (Koike et al., 2007).

Peripheral axonal injury is a clear driver of the transcriptional reprogramming in injured sensory neurons (Cheng et al., 2021; Cho et al., 2013). The results here show that for neurons with large, myelinated axons these transcriptional changes can be induced by demyelination without axonal injury. Further studies are required to determine the mechanism(s) by which Schwann cells dictate the transcriptional state of sensory neurons. Such mechanisms may include an injury signal from Schwann cells to axons provoked by demyelination, such as the release of the toxic metabolite acylcarnitines from Schwann cells (Viader et al., 2013) with demyelination induced mitochondrial dysfunction (Della-Flora Nunes et al., 2021), or signals from myelinating Schwann cells to axons that continuously prevent injury induced transcription reprograming. The axon survival factor nicotinamide mononucleotide adenylyltransferase 2 (NMNAT2) catalyzes the synthesize of nicotinamide adenine dinucleotide (NAD^+^) from nicotinamide mononucleotide (NMN) (Gilley and Coleman, 2010). Axonal injury induces the loss of the axonal NMNAT2, resulting in an impaired NMN to NAD^+^ ratio, which in turn activates the constitutively expressed protein Sterile alpha and Toll/interleukin-1 receptor motif-containing 1 (SARM1) (Figley et al., 2021), resulting in axonal destruction (Coleman and Höke, 2020; Figley and DiAntonio, 2020). Interestingly, maintenance of the mature, myelinating state of Schwann cells also depends on NAD^+^ homeostasis (Sasaki et al., 2018). Nevertheless, a direct link between Schwann cells and axonal factors of survival or destruction has yet to be established. Another mechanism by which Schwann cells affect axons is the metabolic support from glial cells to axons (Beirowski et al., 2014; Fünfschilling et al., 2012; Lee et al., 2012). In line with the different effects of demyelination on specific sensory neurons described here, recent studies revealed that this metabolic support is differentially important for specific neurons (Jha et al., 2020).

We found that PNS demyelination due to Schwann cell ablation does not cause overt widespread axonal loss, which is consistent with a previous study that used the lysolecithin model to induce PNS demyelination (Wallace et al., 2003). Although the authors of the lysolecithin model found that PNS demyelination mainly affects A-fibers, they did not observe induction of ATF3 in the DRG upon PNS demyelination. This can be explained by the inherent difference between the focal demyelination caused by the Lysolecithin to the widespread demyelination caused by genetic ablation of Schwann cells and may also be attribute to the difference in lumber DRGs analyzed. In this study we focused L3-L5 that constitute the sciatic nerve, (Rigaud et al., 2008), whereas L4-L6 were analyzed in the lysolecithin treated sciatic nerves (Wallace et al., 2003).

The PLPCreER^t^ line that was used here (Doerflinger et al., 2003) mediates recombination in both oligodendrocytes and Schwann cells. In addition, recent transcriptomic data suggests that *Plp1* is expressed also in Satellite glial cells (Avraham et al., 2020), suggesting that these cells may be ablated as well. Therefore, new experimental models that will enable separation of CNS demyelination from PNS demyelination, and separation between the effects of Satellite glial cell and Schwann cell ablation are required. Since we were interested here mainly in the transcriptional response of neurons, we used a protocol that we recently developed (Yang et al., 2021) to isolate nuclei from mouse DRGs using iodixanol density gradient centrifugation, which enriches for neuronal nuclei. Further single-RNA-seq studies using an enzymatic-digestion-based protocol, which enriches for non-neuronal cells (Avraham et al., 2020), are required to study the involvement of Satellite glial cells in this model.

Our study has wide implications for the ongoing efforts to identify new therapeutic targets to mitigate PNS neuropathies caused by demyelination, such as CMT1A. Currently, the only available therapy for CMT1A is the experimental drug PXT3003 which is a combination of baclofen, naltrexone, and sorbitol (Attarian et al., 2021). Surprisingly, although CMT1A is caused by demyelination, the beneficial effects of PXT3003 are mainly achieved by targeting axons (Prukop et al., 2020). While the current efforts to identify therapies for PNS demyelination are focused on Schwann cells, aiming to enhance PNS myelination (for example, Fledrich et al., 2018; Zhao et al., 2018), our study suggests that sensory neurons might also be a therapeutic target in demyelinating diseases and that small molecules that target the injury response of sensory neurons, such as camptothecin (Cheng et al., 2021) may be benefitable for PNS demyelination.

## Acknowledgments

This research was supported by the Dr. Miriam and Sheldon G. Adelson Medical Research Foundation (CJW and BP). The authors would like to thank the team of the AMRF Functional Genomics Common Research Resource at the Department of Psychiatry and Neurology, University of California Los Angeles, for their expert help with the sequencing of the Drop-seq libraries. B.P. is supported by NINDS R01NS067550, R01NS109372, R01NS099334 and by the National Multiple Sclerosis Society (RG-1807-32005). B.E. is supported by NINDS R01NS067550. CJW supported by NINDS R35NS105076.

## Author contributions

B.E., C.J.W., and B.P. designed the study. B.E. injected the animals, harvested the tissues, and performed the bulk RNA-seq and TEM analysis. B.L.R. assisted with histological experiments. M.T. provided advice on the usage of the PLP-CreERT;ROSA26-eGFP-DTA mice. W.R. and L.Y. performed the Drop-seq including generation of libraries and bioinformatic analyses. R.K. performed the Next Generation Sequencing. B.E. and B.P. wrote the manuscript, with input from all authors.

## Declaration of interests

The authors declare no competing interests.

## Supplemental Information

### Supplemental Tables

*1. Table S1: Cluster makers for the snRNAseq data*.

*2. Table S2-S4: (S2) Number of cells by annotation, (S3) Number of cells by cell type, (S4) Number of injured cells by cell type*.

*3. Table S5: induced genes in the injured cluster compared to all other naÏve clusters*.

### Supplemental Figures

**Figure S1:**
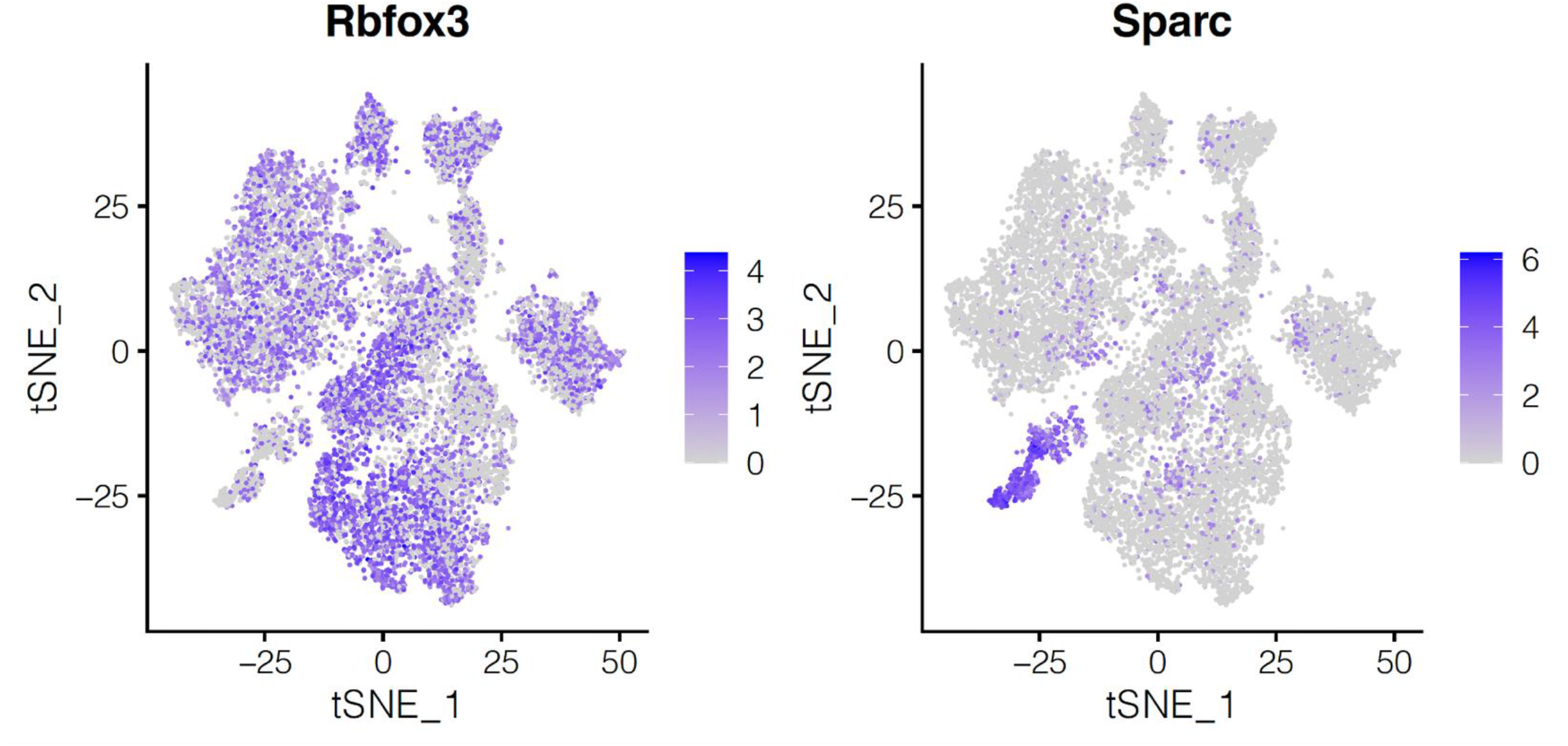
tSNE plots of the neuronal cluster expressing *Rbfox3*, and non-neuronal clusters expressing *Sparc*. Nuclei are colored by their log2 expression of the neuronal marker gene *Rbfox3* (top) and non-neuronal marker gene, *Sparc* (bottom).

**Figure S2:**
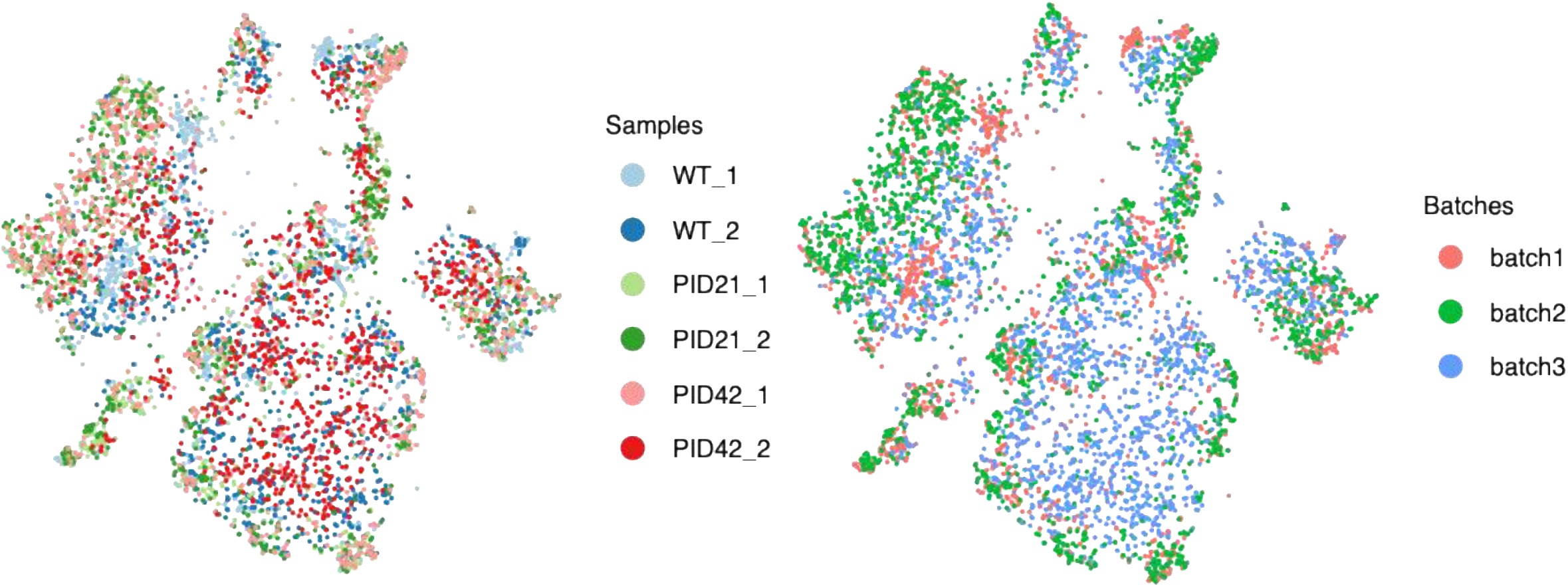
tSNE plot of the nuclei derived from three biological replicates of the experimental conditions. Nuclei are colored by their biological replicate. Each cluster has a mix of nuclei from each replicate, suggesting there are minimal if any batch effects between replicates.

**Figure S3:**
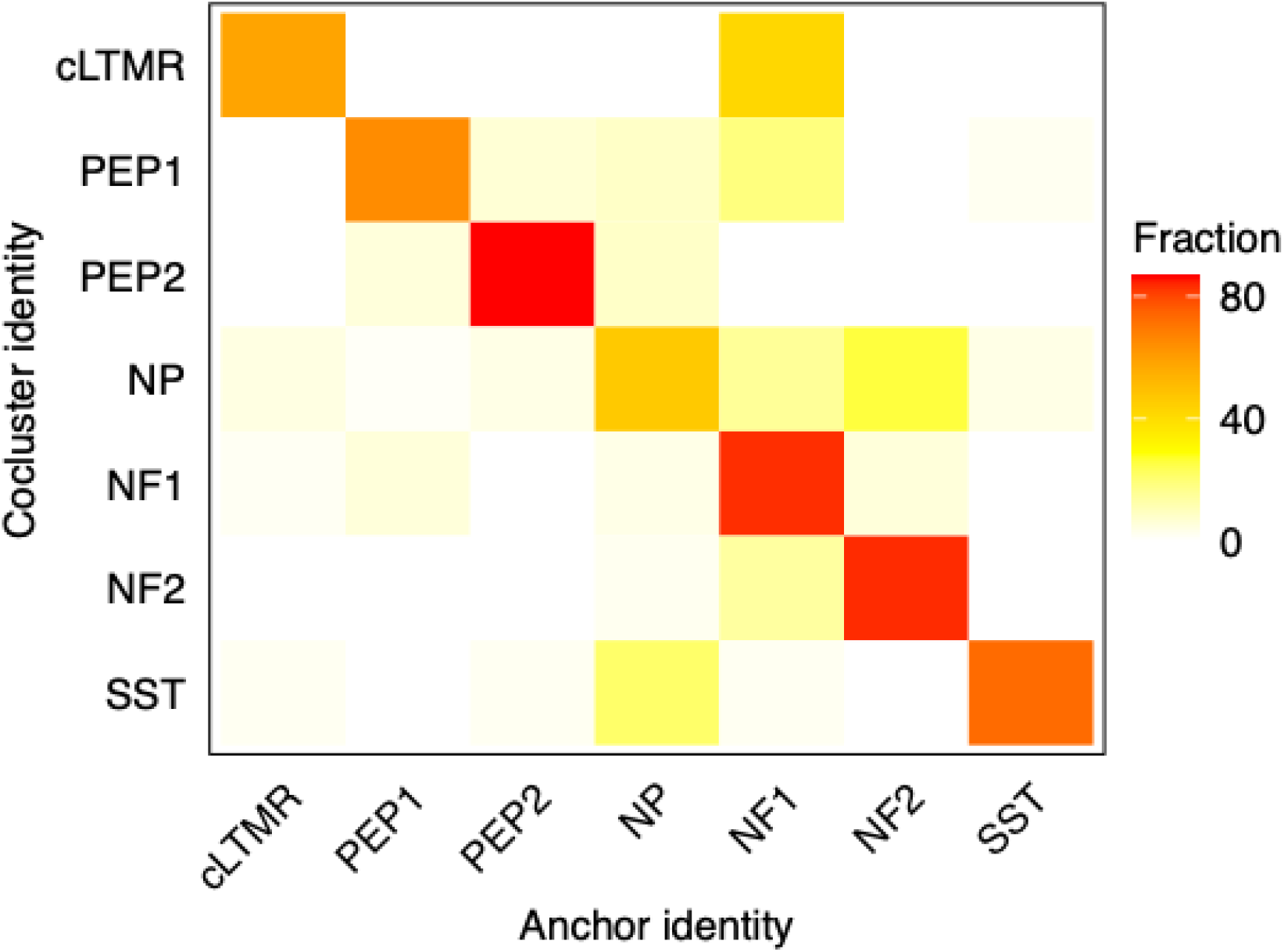
Classifications of injured nuclei. Overlap of cell type classifications of injured nuclei between coclustering with Spinal Nerve Transection (from Figure 4A) and anchoring to the uninjured clusters (See methods). The fraction of injured nuclei within the initial cell type assignment that is assigned to each cell type after anchoring is displayed.

**Fig S4.**
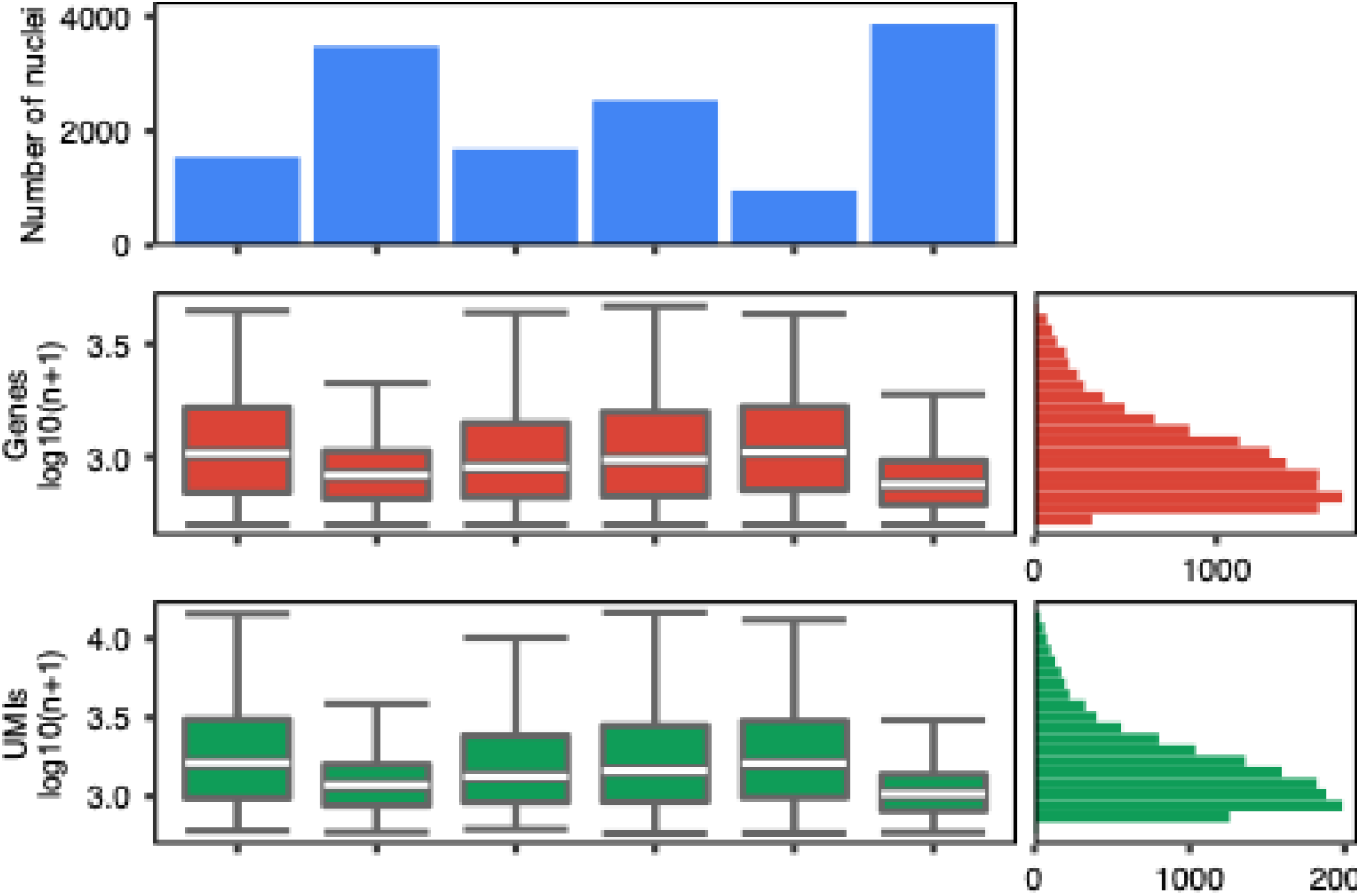
snRNA-seq metrics for all sequencing samples. Number of nuclei from each library with > 500 unique genes (top box). Box plots display number of genes per nucleus (log10 transformed, middle row), and unique molecular identifiers (UMI) per nucleus (log10 transformed, bottom row). Boxes indicate quartiles and whiskers are 1.5-times the interquartile range (Q1-Q3). The median is a white line inside each box. The distribution is aggregated across all samples and displayed on the horizontal histogram.

## STAR Methods

### Key Resources Table

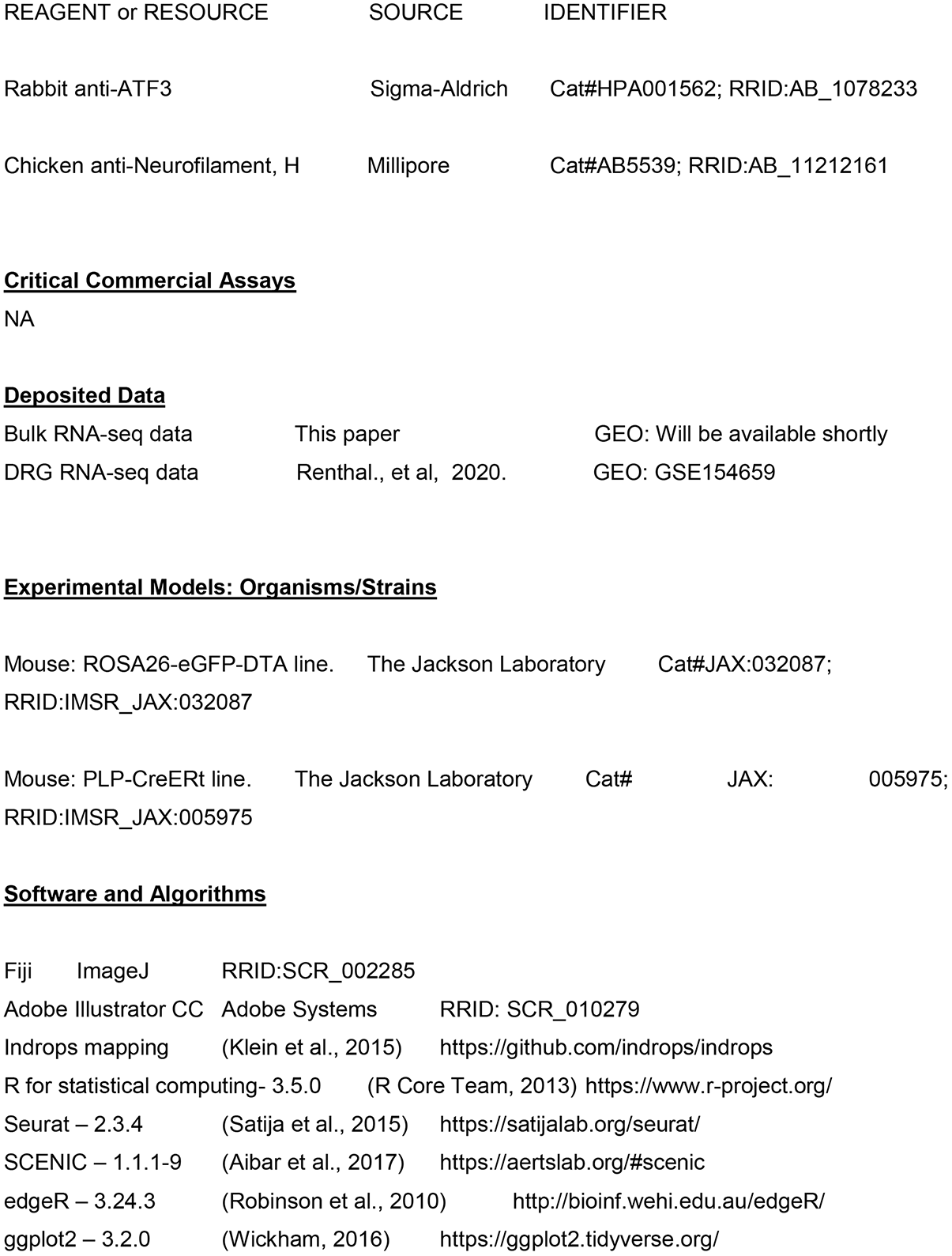
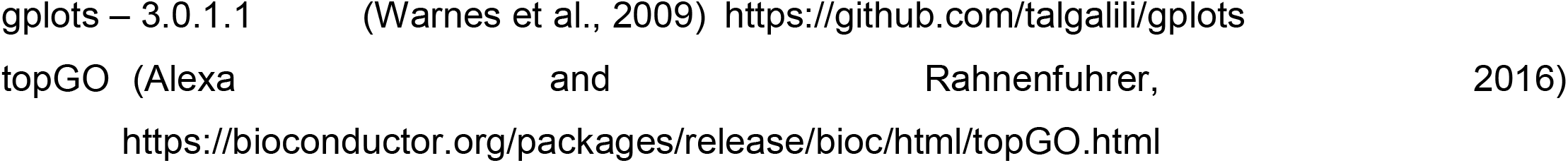

## Resource Availability

### Lead Contact

Further information and requests for resources and reagents should be directed to and will be fulfilled by the Lead Contact, Benayahu Elbaz.

### Materials Availability

The PLP-CreERt-ROSA26-eGFP-DTA mouse model used in this study is available from Brian Popko upon request.

### Data and Code Availability

Raw and processed data were deposited within the Gene Expression Omnibus (GEO) repository (https://www.ncbi.nlm.nih.gov/geo) and will be available shortly. Custom R scripts are available upon request from William Renthal.

### Experimental Model and Subject Details

### Mice

Plp-CreERt;ROSA26-eGFP-DTA mice and Cre negative ROSA26-eGFP-DTA mice (used as control) were generated by crossing the ROSA26-eGFP-DT-A mice (Cat#JAX:032087; RRID:IMSR_JAX:032087) with the PLP-CreERt mice (Doerflinger et al., 2003) (Cat# JAX: 005975; RRID:IMSR_JAX:005975). All animal experiments were conducted according to institutional animal care and safety guidelines.

## Method Details

### Dissection Procedures

Tamoxifen treated mice were euthanized by CO2 asphyxiation and decapitation. Lumbar L3-L5 ganglia were collected at various 21 days and 42 days post injection. Ganglia from 3-5 mice per sample were immediately frozen on dry ice, then pooled for subsequent snRNA-seq profiling or histology. There were 2-3 biological replicates of each pooled condition.

### Tamoxifen injections

0.8 mg/day 4-hydroxytamoxifen (H-6278, Sigma) was injected for 3 consecutive days for females, and for 4 consecutive days for males, based on protocol developed for the Plp-CreERt;ROSA26-eGFP-DTA mice (Traka, 2019).

### Single-nuclei isolation from mouse DRG

Single-nuclei suspensions of lumbar DRGs from treated mice were collected using a recently described protocol (Yang et al., 2021). This method increases the isolation of neuronal nuclei compared to commonly used nuclear extraction protocols (Mo et al., 2015; Renthal et al., 2018). Briefly, DRGs were removed from dry ice and placed into homogenization buffer (0.25 M sucrose, 25 mM KCl, 5 mM MgCl2, 20 mM tricine-KOH, pH 7.8, 1 mM DTT, 5 μg/mL actinomycin). After a brief incubation on ice, the samples were briefly homogenized using a Tissue-Tearor and transferred to a Dounce homogenizer for an additional ten strokes with a tight pestle in a total volume of 5mL homogenization buffer. After ten strokes with a tight pestle, a 5% IGEPAL (Sigma) solution was added to a final concentration of 0.32% and five additional strokes with the tight pestle were formed. The tissue homogenate was then passed through a 20-μm filter, and diluted 1:1 with OptiPrep (Sigma) and layered onto an OptiPrep gradient. After ultracentrifugation, nuclei were collected between the 30-40% OptiPrep layers. This layer contains DRG nuclei as well as some membrane fragments likely from Schwann cells that have the same density as nuclei. We diluted this layer in 30% OptiPrep to a final concentration of 80,000 - 90,000 nuclei /mL for loading into the inDrops microfluidic device. All buffers and gradient solutions for nuclei extraction contained 0.01U/ul RNase inhibitor (Promega) and 0.04% BSA.

### Single-nucleus RNA sequencing (inDrops)

Single-nuclei suspensions were encapsulated into droplets and the RNA in each droplet was reverse transcribed using a unique oligonucleotide barcode for each nucleus as described previously (Renthal et al., 2020). Nuclei encapsulation was performed in a blinded fashion and the order of sample processing was randomized. The total number of droplets collected per sample varied based on available reagents and line integrity. After encapsulation, each sample was divided into pools of approximately 3,000 droplets and library preparation was performed on each pool of droplets. Libraries were sequenced on an Illumina Nextseq 500 to a depth of 500 million reads per ∼30,000 droplets collected, resulting in at least 5 reads per UMI on average per sample. Sequencing data was processed and mapped to the mouse genome using the pipeline described in https://github.com/indrops/indrops (Klein et al., 2015). Counts tables from each library were then combined and processed as described below. The metrics for all sequencing samples is presented in Figure S4.

### Initial quality control, clustering and visualization of snRNA-seq

To be included for analysis, nuclei were required to contain counts of greater than 500 unique genes, fewer than 15,000 total UMI, and fewer than 5% of the counts deriving from mitochondrial genes. We used the Seurat package (version 2.3.4) in R (R Core Team, 2013) to perform clustering of these nuclei as previously described (Satija et al., 2015). Raw counts were scaled to 10,000 transcripts per nucleus to control the sequencing depth between nuclei. Counts were centered and scaled for each gene. The effects of total UMI and percent of mitochondrial genes in each nucleus were regressed out using a linear model in the Scaledata() function. Highly variable genes were identified using the MeanVarPlot() with default settings. The top 20 principal components were retrieved with the RunPCA() function using default parameters. Nuclei clustering was performed using FindClusters() based on the top 20 principal components, with resolution at 1 for the initial clustering of all nuclei and the sub-clustering of non-neuronal nuclei except where otherwise specified. For dimension reduction and visualization, tSNE coordinates were calculated in the PCA space by using the implemented function runTSNE() in Seurat.

### Doublet identification and classification of DRG subtypes

Marker genes for each cluster were identified by running FindAllMarkers() in Seurat comparing nuclei in one cluster to all other nuclei. Doublet or low-quality clusters were identified as clusters which either are significantly enriched for at least two mitochondrial genes (Log2FC > 0.5, FDR < 0.05), or have no significantly enriched cluster marker genes (FDR < 0.05, log2FC > 1) other than Rgs11. Nuclei in those clusters were excluded from the dataset. The remaining nuclei were clustered again, and marker gene analysis was run as described above. Significant enrichment (FDR < 0.05, log2FC > 0.5) of known subtype marker genes (peptidergic nociceptors (PEP) = Tac1, non-peptidergic nociceptors (NP) = Cd55, pruriceptors (SST) = Sst, cLTMR = Fam19a4, A-LTMR (NF) = Nefh, Satglia = Apoe, Schwann cells = Mpz) within a cluster of nuclei compared to all other nuclei was used to assign the subtype to each of the naive cluster. In our samples Cadps2+ Aδ-LTMRs nuclei were rare and were not classified as a separate cluster.

### Identification of injured clusters and subtype classification of of injured nuclei

Clusters of cells that are in a transcriptionally injured state were identified as the clusters that are significantly enriched of *Atf3* (FDR < 0.05, log2FC > 1) compared to all other clusters. To identify the subtypes for injured nuclei, we took two different approaches. First, All injured nuclei from the demyelination model were co-clustered with injured nuclei from spinal nerve transection (SpNT) as described previously (Renthal et al., 2020), and clusters were identified. Having classified SpNT nuclei from the previous study, we were able to project those neuronal subtypes onto the injured state nuclei from the demyelination model. For each cluster, the subtype pf the most abundant previously-classified injured SpNT was projected onto all unclassified nuclei in that cluster (95 ± 3% of previously-classified injured SpNT nuclei in each cluster share the same subtype label). Cell type labels classified using this approach were used for all analysis unless otherwise specified. As an independent approach, we also anchored all injured nuclei from the demyelination model to all other nuclei that are in a transcriptionally uninjured state from the same model. FindTransferAnchors(reduction = ‘‘cca’’) in Seurat was then used to identify anchors between injured and mouse data. TransferData() was used to transfer uninjured subtype labels to each nucleus in the injured state.

### Differential expression analysis

To identify genes that are differentially expressed in injured NF2 nuclei, differential expression analysis was done with edgeR (version 3.24.3) as we recently described (Renthal et al., 2020). Briefly, edgeR uses the raw counts as input, and genes detected in fewer than 5% of nuclei selected for each comparison were excluded from analysis. Counts within each nucleus were normalized by the trimmed mean of M-values (TMM) method to adjust for total RNA differences between nuclei. Dispersion was estimated by fitting a quasi-likelihood negative binomial generalized log-linear model (glmQLFit) with the conditions being analyzed. The QL F-test was used to determine statistical significance between differentially expressed genes in the experimental and control group. The experimental group was identified as 166 injured NF2 nuclei from PID21 samples. The control group was selected by randomly sampling the same number of NF2 nuclei from no disease samples. Sampling of control group and differential expression analysis was repeated ten times and median Log2FC and FDR for each gene was reported.

### Immunohistochemistry

L3-L5 DRGs were harvested from treated mice perfused with cold PBS followed by cold 4% PFA. The perfused tissues were fixed in 4% PFA for 1hr at 4°C and cryoprotected with 30% sucrose in PBS overnight. DRGs were sectioned into 12μm sections, which were blocked and permeabilized with 1% Triton X-100 in blocking buffer for 1hr at 25°C. Sections were incubated with rabbit polyclonal antibody against ATF3 ([1:1000]; Santa Cruz Biotech; Cat#sc-188; RRID:AB_2258513) and chicken polyclonal antibody against NF200 ([1:2000]; Millipore; AB5539; RRID:AB_11212161) at 4°C overnight and then incubated with Alexa Fluor 568 goat antibody against rabbit IgG and Alexa Fluor 488 goat antibody against chicken IgG for 1hr at 25°C. Images were acquired using a Slide Scanner microscope.

### Data Visualization

Plots were generated using R version 3.5.0 with ggplot2 package (version 3.2.0) (Wickham, 2016). Heatmaps were generated using gplots package (version 3.0.1.1) (Warnes et al., 2009). Figures were made using Adobe Illustrator (Adobe Systems; RRID: SCR_010279).

## Quantification and Statistical Analysis

### Statistical analysis

Statistical analyses including the number of animals or cells (n) and P values for each experiment are noted in the figure legends. Statistics were performed using R version 3.5.0. Hypergeometric tests were used to test the significance of overlap between two gene sets. It was conducted by calling the phyper() function in R version 3.5.0.

## Notes

### Competing Interest Statement

The authors have declared no competing interest.

## Bibliography

Abe, N., and Cavalli, V. (2008). Nerve injury signaling. Curr. Opin. Neurobiol. 18, 276–283.

Aibar, S., González-Blas, C.B., Moerman, T., Huynh-Thu, V.A., Imrichova, H., Hulselmans, G., Rambow, F., Marine, J.-C., Geurts, P., Aerts, J., et al. (2017). SCENIC: single-cell regulatory network inference and clustering. Nat. Methods 14, 1083–1086.

Alexa, A., and Rahnenfuhrer, J. (2016). topGO: Enrichment analysis for Gene Ontology. R package version 2.28. 0. Cranio.

Arthur-Farraj, P.J., Latouche, M., Wilton, D.K., Quintes, S., Chabrol, E., Banerjee, A., Woodhoo, A., Jenkins, B., Rahman, M., Turmaine, M., et al. (2012). c-Jun reprograms Schwann cells of injured nerves to generate a repair cell essential for regeneration. Neuron 75, 633–647.

Attarian, S., Young, P., Brannagan, T.H., Adams, D., Van Damme, P., Thomas, F.P., Casanovas, C., Tard, C., Walter, M.C., Péréon, Y., et al. (2021). A double-blind, placebo-controlled, randomized trial of PXT3003 for the treatment of Charcot-Marie-Tooth type 1A. Orphanet J Rare Dis 16, 433.

Avraham, O., Deng, P.-Y., Jones, S., Kuruvilla, R., Semenkovich, C.F., Klyachko, V.A., and Cavalli, V. (2020). Satellite glial cells promote regenerative growth in sensory neurons. Nat. Commun. 11, 4891.

Beirowski, B., Babetto, E., Golden, J.P., Chen, Y.-J., Yang, K., Gross, R.W., Patti, G.J., and Milbrandt, J. (2014). Metabolic regulator LKB1 is crucial for Schwann cell-mediated axon maintenance. Nat. Neurosci. 17, 1351–1361.

Benito, C., Davis, C.M., Gomez-Sanchez, J.A., Turmaine, M., Meijer, D., Poli, V., Mirsky, R., and Jessen, K.R. (2017). STAT3 Controls the Long-Term Survival and Phenotype of Repair Schwann Cells during Nerve Regeneration. J. Neurosci. 37, 4255–4269.

Berger, P., Niemann, A., and Suter, U. (2006). Schwann cells and the pathogenesis of inherited motor and sensory neuropathies (Charcot-Marie-Tooth disease). Glia 54, 243–257.

Catenaccio, A., Llavero Hurtado, M., Diaz, P., Lamont, D.J., Wishart, T.M., and Court, F.A. (2017). Molecular analysis of axonal-intrinsic and glial-associated co-regulation of axon degeneration. Cell Death Dis. 8, e3166.

Chandran, V., Coppola, G., Nawabi, H., Omura, T., Versano, R., Huebner, E.A., Zhang, A., Costigan, M., Yekkirala, A., Barrett, L., et al. (2016). A Systems-Level Analysis of the Peripheral Nerve Intrinsic Axonal Growth Program. Neuron 89, 956–970.

Cheng, Y.-C., Snavely, A., Barrett, L.B., Zhang, X., Herman, C., Frost, D.J., Riva, P., Tochitsky, I., Kawaguchi, R., Singh, B., et al. (2021). Topoisomerase I inhibition and peripheral nerve injury induce DNA breaks and ATF3-associated axon regeneration in sensory neurons. Cell Rep. 36, 109666.

Cho, Y., Sloutsky, R., Naegle, K.M., and Cavalli, V. (2013). Injury-induced HDAC5 nuclear export is essential for axon regeneration. Cell 155, 894–908.

Coleman, M.P., and Höke, A. (2020). Programmed axon degeneration: from mouse to mechanism to medicine. Nat. Rev. Neurosci. 21, 183–196.

Corfas, G., Velardez, M.O., Ko, C.-P., Ratner, N., and Peles, E. (2004). Mechanisms and roles of axon-Schwann cell interactions. J. Neurosci. 24, 9250–9260.

Della-Flora Nunes, G., Wilson, E.R., Hurley, E., He, B., O’Malley, B.W., Poitelon, Y., Wrabetz, L., and Feltri, M.L. (2021). Activation of mTORC1 and c-Jun by Prohibitin1 loss in Schwann cells may link mitochondrial dysfunction to demyelination. Elife 10.

Doerflinger, N.H., Macklin, W.B., and Popko, B. (2003). Inducible site-specific recombination in myelinating cells. Genesis 35, 63–72.

Figley, M.D., and DiAntonio, A. (2020). The SARM1 axon degeneration pathway: control of the NAD+ metabolome regulates axon survival in health and disease. Curr. Opin. Neurobiol. 63, 59–66.

Figley, M.D., Gu, W., Nanson, J.D., Shi, Y., Sasaki, Y., Cunnea, K., Malde, A.K., Jia, X., Luo, Z., Saikot, F.K., et al. (2021). SARM1 is a metabolic sensor activated by an increased NMN/NAD+ ratio to trigger axon degeneration. Neuron 109, 1118–1136.e11.

Fledrich, R., Abdelaal, T., Rasch, L., Bansal, V., Schütza, V., Brügger, B., Lüchtenborg, C., Prukop, T., Stenzel, J., Rahman, R.U., et al. (2018). Targeting myelin lipid metabolism as a potential therapeutic strategy in a model of CMT1A neuropathy. Nat. Commun. 9, 3025.

Fünfschilling, U., Supplie, L.M., Mahad, D., Boretius, S., Saab, A.S., Edgar, J., Brinkmann, B.G., Kassmann, C.M., Tzvetanova, I.D., Möbius, W., et al. (2012). Glycolytic oligodendrocytes maintain myelin and long-term axonal integrity. Nature 485, 517–521.

Gerber, D., Ghidinelli, M., Tinelli, E., Somandin, C., Gerber, J., Pereira, J.A., Ommer, A., Figlia, G., Miehe, M., Nägeli, L.G., et al. (2019). Schwann cells, but not Oligodendrocytes, Depend Strictly on Dynamin 2 Function. Elife 8.

Gilley, J., and Coleman, M.P. (2010). Endogenous Nmnat2 is an essential survival factor for maintenance of healthy axons. PLoS Biol. 8, e1000300.

Gonçalves, N.P., Vægter, C.B., Andersen, H., Østergaard, L., Calcutt, N.A., and Jensen, T.S. (2017). Schwann cell interactions with axons and microvessels in diabetic neuropathy. Nat. Rev. Neurol. 13, 135–147.

Imai, S., Koyanagi, M., Azimi, Z., Nakazato, Y., Matsumoto, M., Ogihara, T., Yonezawa, A., Omura, T., Nakagawa, S., Wakatsuki, S., et al. (2017). Taxanes and platinum derivatives impair Schwann cells via distinct mechanisms. Sci. Rep. 7, 5947.

Jessen, K.R., and Mirsky, R. (2016). The repair Schwann cell and its function in regenerating nerves. J. Physiol. (Lond.) 594, 3521–3531.

Jessen, K.R., and Mirsky, R. (2019). The success and failure of the schwann cell response to nerve injury. Front. Cell Neurosci. 13, 33.

Jessen, K.R., Mirsky, R., and Lloyd, A.C. (2015). Schwann cells: development and role in nerve repair. Cold Spring Harb. Perspect. Biol. 7, a020487.

Jha, M.K., Lee, Y., Russell, K.A., Yang, F., Dastgheyb, R.M., Deme, P., Ament, X.H., Chen, W., Liu, Y., Guan, Y., et al. (2020). Monocarboxylate transporter 1 in Schwann cells contributes to maintenance of sensory nerve myelination during aging. Glia 68, 161–177.

Klein, A.M., Mazutis, L., Akartuna, I., Tallapragada, N., Veres, A., Li, V., Peshkin, L., Weitz, D.A., and Kirschner, M.W. (2015). Droplet barcoding for single-cell transcriptomics applied to embryonic stem cells. Cell 161, 1187–1201.

Koike, H., Iijima, M., Mori, K., Yamamoto, M., Hattori, N., Katsuno, M., Tanaka, F., Watanabe, H., Doyu, M., Yoshikawa, H., et al. (2007). Nonmyelinating Schwann cell involvement with well-preserved unmyelinated axons in Charcot-Marie-Tooth disease type 1A. J. Neuropathol. Exp. Neurol. 66, 1027–1036.

Lee, Y., Morrison, B.M., Li, Y., Lengacher, S., Farah, M.H., Hoffman, P.N., Liu, Y., Tsingalia, A., Jin, L., Zhang, P.-W., et al. (2012). Oligodendroglia metabolically support axons and contribute to neurodegeneration. Nature 487, 443–448.

Ma, K.H., and Svaren, J. (2018). Epigenetic control of schwann cells. Neuroscientist 24, 627–638.

Michailov, G.V., Sereda, M.W., Brinkmann, B.G., Fischer, T.M., Haug, B., Birchmeier, C., Role, L., Lai, C., Schwab, M.H., and Nave, K.-A. (2004). Axonal neuregulin-1 regulates myelin sheath thickness. Science 304, 700–703.

Mindos, T., Dun, X.-P., North, K., Doddrell, R.D.S., Schulz, A., Edwards, P., Russell, J., Gray, B., Roberts, S.L., Shivane, A., et al. (2017). Merlin controls the repair capacity of Schwann cells after injury by regulating Hippo/YAP activity. J. Cell Biol. 216, 495–510.

Mo, A., Mukamel, E.A., Davis, F.P., Luo, C., Henry, G.L., Picard, S., Urich, M.A., Nery, J.R., Sejnowski, T.J., Lister, R., et al. (2015). Epigenomic signatures of neuronal diversity in the mammalian brain. Neuron 86, 1369–1384.

Nguyen, M.Q., Le Pichon, C.E., and Ryba, N. (2019). Stereotyped transcriptomic transformation of somatosensory neurons in response to injury. Elife 8.

Prukop, T., Wernick, S., Boussicault, L., Ewers, D., Jäger, K., Adam, J., Winter, L., Quintes, S., Linhoff, L., Barrantes-Freer, A., et al. (2020). Synergistic PXT3003 therapy uncouples neuromuscular function from dysmyelination in male Charcot-Marie-Tooth disease type 1A (CMT1A) rats. J. Neurosci. Res. 98, 1933–1952.

R Core Team (2013). R: A language and environment for statistical computing (Vienna, Austria: R Foundation for Statistical Computing).

Renthal, W., Boxer, L.D., Hrvatin, S., Li, E., Silberfeld, A., Nagy, M.A., Griffith, E.C., Vierbuchen, T., and Greenberg, M.E. (2018). Characterization of human mosaic Rett syndrome brain tissue by single-nucleus RNA sequencing. Nat. Neurosci. 21, 1670–1679.

Renthal, W., Tochitsky, I., Yang, L., Cheng, Y.-C., Li, E., Kawaguchi, R., Geschwind, D.H., and Woolf, C.J. (2020). Transcriptional Reprogramming of Distinct Peripheral Sensory Neuron Subtypes after Axonal Injury. Neuron 108, 128–144.e9.

Rigaud, M., Gemes, G., Barabas, M.-E., Chernoff, D.I., Abram, S.E., Stucky, C.L., and Hogan, Q.H. (2008). Species and strain differences in rodent sciatic nerve anatomy: implications for studies of neuropathic pain. Pain 136, 188–201.

Robinson, M.D., McCarthy, D.J., and Smyth, G.K. (2010). edgeR: a Bioconductor package for differential expression analysis of digital gene expression data. Bioinformatics 26, 139–140.

Sasaki, Y., Hackett, A.R., Kim, S., Strickland, A., and Milbrandt, J. (2018). Dysregulation of NAD+ metabolism induces a schwann cell dedifferentiation program. J. Neurosci. 38, 6546–6562.

Satija, R., Farrell, J.A., Gennert, D., Schier, A.F., and Regev, A. (2015). Spatial reconstruction of single-cell gene expression data. Nat. Biotechnol. 33, 495–502.

Schmalbruch, H. (1986). Fiber composition of the rat sciatic nerve. Anat Rec 215, 71–81.

Sergeeva, E.G., Rosenberg, P.A., and Benowitz, L.I. (2021). Non-Cell-Autonomous Regulation of Optic Nerve Regeneration by Amacrine Cells. Front. Cell Neurosci. 15, 666798.

Sharma, N., Flaherty, K., Lezgiyeva, K., Wagner, D.E., Klein, A.M., and Ginty, D.D. (2020). The emergence of transcriptional identity in somatosensory neurons. Nature 577, 392–398.

Taveggia, C., Zanazzi, G., Petrylak, A., Yano, H., Rosenbluth, J., Einheber, S., Xu, X., Esper, R.M., Loeb, J.A., Shrager, P., et al. (2005). Neuregulin-1 type III determines the ensheathment fate of axons. Neuron 47, 681–694.

Traka, M. (2019). The DTA mouse model for oligodendrocyte ablation and CNS demyelination. Methods Mol. Biol. 1936, 295–310.

Traka, M., Arasi, K., Avila, R.L., Podojil, J.R., Christakos, A., Miller, S.D., Soliven, B., and Popko, B. (2010). A genetic mouse model of adult-onset, pervasive central nervous system demyelination with robust remyelination. Brain 133, 3017–3029.

Traka, M., Podojil, J.R., McCarthy, D.P., Miller, S.D., and Popko, B. (2016). Oligodendrocyte death results in immune-mediated CNS demyelination. Nat. Neurosci. 19, 65–74.

Usoskin, D., Furlan, A., Islam, S., Abdo, H., Lönnerberg, P., Lou, D., Hjerling-Leffler, J., Haeggström, J., Kharchenko, O., Kharchenko, P.V., et al. (2015). Unbiased classification of sensory neuron types by large-scale single-cell RNA sequencing. Nat. Neurosci. 18, 145–153.

Vaquié, A., Sauvain, A., Duman, M., Nocera, G., Egger, B., Meyenhofer, F., Falquet, L., Bartesaghi, L., Chrast, R., Lamy, C.M., et al. (2019). Injured axons instruct schwann cells to build constricting actin spheres to accelerate axonal disintegration. Cell Rep. 27, 3152–3166.e7.

Viader, A., Sasaki, Y., Kim, S., Strickland, A., Workman, C.S., Yang, K., Gross, R.W., and Milbrandt, J. (2013). Aberrant Schwann cell lipid metabolism linked to mitochondrial deficits leads to axon degeneration and neuropathy. Neuron 77, 886–898.

Wallace, V.C.J., Cottrell, D.F., Brophy, P.J., and Fleetwood-Walker, S.M. (2003). Focal lysolecithin-induced demyelination of peripheral afferents results in neuropathic pain behavior that is attenuated by cannabinoids. J. Neurosci. 23, 3221–3233.

Warnes, G.R., Bolker, B., Bonebakker, L., Gentleman, R., Huber, W., Liaw, A., Lumley, T., Maechler, M., Magnusson, A., Moeller, S., et al. (2009). gplots: Various R programming tools for plotting data. R Package Version 2.

Wickham, H. (2016). ggplot2 - Elegant Graphics for Data Analysis (New York, NY: Springer-Verlag New York).

Yang, L., Tochitsky, I., Woolf, C.J., and Renthal, W. (2021). Isolation of Nuclei from Mouse Dorsal Root Ganglia for Single-nucleus Genomics. Bio Protoc 11, e4102.

Zeisel, A., Hochgerner, H., Lönnerberg, P., Johnsson, A., Memic, F., van der Zwan, J., Häring, M., Braun, E., Borm, L.E., La Manno, G., et al. (2018). Molecular architecture of the mouse nervous system. Cell 174, 999–1014.e22.

Zhang, Y., Williams, P.R., Jacobi, A., Wang, C., Goel, A., Hirano, A.A., Brecha, N.C., Kerschensteiner, D., and He, Z. (2019). Elevating growth factor responsiveness and axon regeneration by modulating presynaptic inputs. Neuron 103, 39–51.e5.

Zhao, H.T., Damle, S., Ikeda-Lee, K., Kuntz, S., Li, J., Mohan, A., Kim, A., Hung, G., Scheideler, M.A., Scherer, S.S., et al. (2018). PMP22 antisense oligonucleotides reverse Charcot-Marie-Tooth disease type 1A features in rodent models. J. Clin. Invest. 128, 359–368.

Zheng, Y., Liu, P., Bai, L., Trimmer, J.S., Bean, B.P., and Ginty, D.D. (2019). Deep sequencing of somatosensory neurons reveals molecular determinants of intrinsic physiological properties. Neuron 103, 598–616.e7.

